# The reactivation of reprogramming factors within human blastocysts by using ATP contribute to human blastocyst development

**DOI:** 10.1101/111708

**Authors:** Takashi Takemura, Midori Okabe

## Abstract

Scientists worldwide have been unable to replicate the stimulus-triggered acquisition of pluripotency (STAP) cells and/or the STAP phenomenon. However, investigations into STAP cells and/or the STAP phenomenon by RIKEN CDB in Japan found that ATP (adenosine 5’-triphosphate disodium salt hydrate) can upregulate Oct3/4 (POU5F1: POU domain, class 5, transcription factor 1) and Nanog mRNA expression in mouse hepatocytes.

On the other hand, no studies have investigated whether ATP can contribute to human blastocyst development. Here we show the reactivation of reprogramming factors within human blastocysts by appropriate ATP treatment (1 mM for 2 days) can contribute to human blastocyst development.

In conclusion, although ATP treatment could not replicate STAP cells and/or the STAP phenomenon by scientists worldwide, appropriate ATP treatment (1 mM for 2 days) in cultured human blastocysts with totipotency would be helpful for infertility women.

## Introduction

An investigation have shown that ATP (adenosine 5’-triphosphate disodium salt hydrate) can upregulate Oct3/4 (also known as POU5F1: POU domain, class 5, transcription factor 1) and Nanog mRNA expression in mouse hepatocytes [1].

Oct3/4 is an important regulatory factor during early embryonic development [2]. Furthermore, Nanog is a transcription factor expressed in the inner cell mass (ICM) within blastocysts [3].

Therefore, ATP treatment of human blastocysts may promote human blastocyst development. In addition, Oct3/4 and Nanog are known as reprogramming factors to generate induced pluripotent stem (iPS) cells [4, 5].

However, no studies have investigated whether ATP can contribute to human blastocyst development. Therefore, in the present study, we investigated the influence of ATP on human blastocyst development.

## Materials and Methods

### Study design

The present study was an experimental study involving 105 patients (median age: 34.2 years) in a previous study [6] with infertility at our institute.

### Discarded human blastocysts and the study protocol

Discarded human blastocysts (Day 5) were collected from patients in a previous study [6]. The influence of ATP on human blastocyst development was studied (untreated group as control group, n=50; ATP-exposed group, n=55) based on the Gardner and Schoolcraft scoring system [7]. Human blastocyst development was scored by blastocoel stage (one to six, highest score is six), ICM grade (highest score A, followed by B and C) and trophectoderm (TE) grade (highest score A, followed by B and C). Higher scores were considered improvements. Two clinical embryologists evaluated the human blastocyst development. Diluted ATP solution was prepared by diluting ATP (Sigma, USA) in distilled water to a final concentration of 200 mM based on the previous protocol [1] or a final concentration of 1 mM.

In the ATP-exposed group, human blastocysts (Day 5) were treated with either 200 mM (n=10) or 1 mM (n=45) of ATP (Sigma, USA) for 2 days and were cultured in Sydney IVF Blastocyst Medium (Cook, USA).

### Quantitative RT-PCR

By using the discarded human blastocysts [6], the relative mRNA expression level of Oct3/4 and Nanog within the human blastocysts were analyzed in the untreated group and ATP-exposed groups using quantitative reverse transcription polymerase chain reaction (qRT-PCR) with an ABI StepOne Real-Time PCR System (Applied Biosystems, USA) according to previously described methods [8]. Furthermore, the PCR primers for Oct3/4 and Nanog were based on previously described primer sequences [8].

### Institutional Review Board (IRB) approval

This study was approved by the IRB of Reproductive Medicine Institute, Japan, Tokyo. Furthermore, the patients provided informed consent.

### Statistical analyses

Statistical analyses were performed using Dr. SPSS II for Windows (SPSS Japan, Inc., Tokyo), and significance was defined as p<0.05. Fisher's exact test was used in the statistical analyses.

## Results

### Human blastocysts in the ATP-exposed group

In the ATP-exposed group, all human blastocysts (Day 5) that were treated with ATP (200 mM for 2 days, n = 10; Sigma, USA) were bursted. Conversely, all human blastocysts (Day 5) that were treated with ATP (1 mM for 2 days, n = 45; Sigma, USA) survived.

### ATP treatment (1 mM for 2 days, n = 45) improves the ICM and TE grade within human blastocysts (Day 5) compared with the untreated group as the control group

Fisher's exact test showed that the ICM grade was significantly higher in the ATP-exposed group (p < 0.001) than the untreated group (31/45 [68.9%] vs. 0/50 [0%], respectively) (Table 1).

**Table 1:**
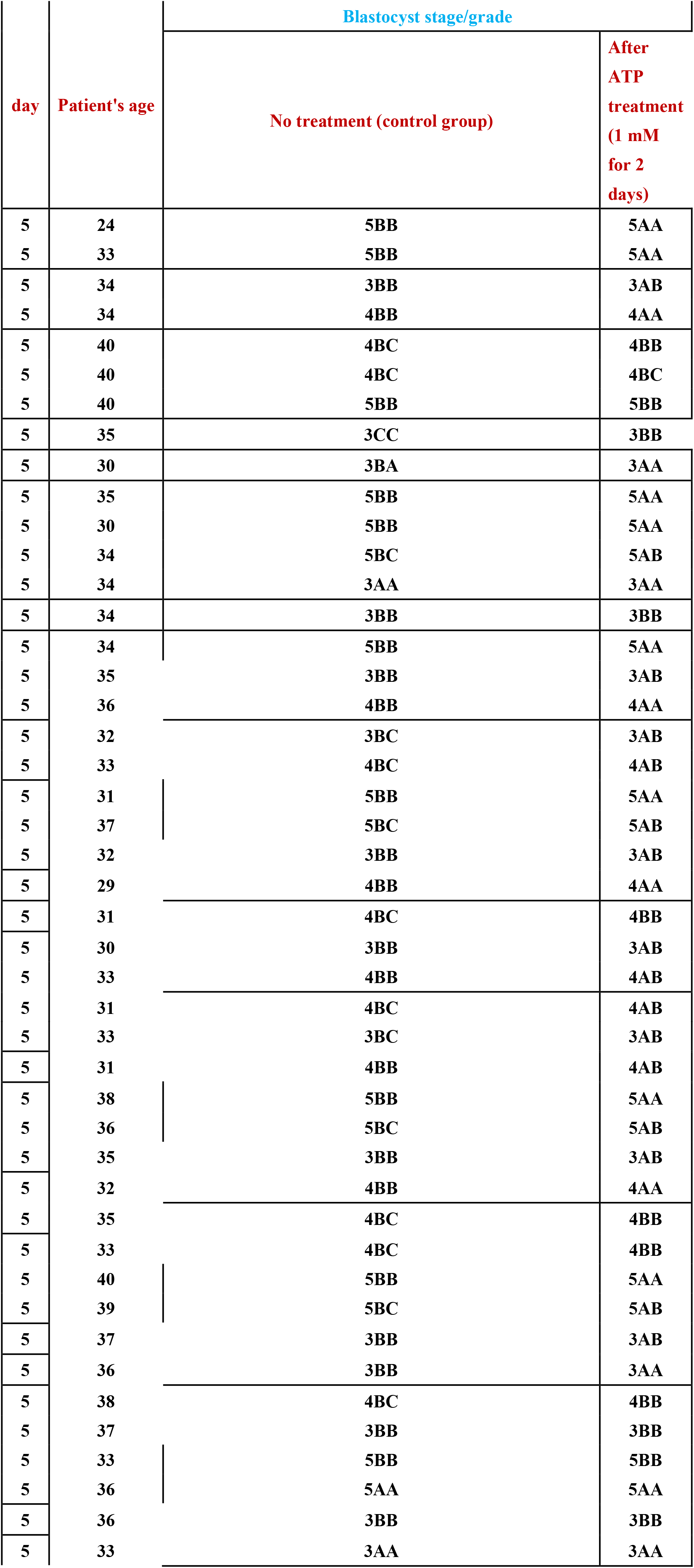
Changes in the ICM grade and TE grade within human blastocysts (Day 5) treated with ATP (1 mM for 2 days, n= 45).

Furthermore, Fisher's exact test showed that the TE grade was significantly higher in the ATP-exposed group (p < 0.001) than the untreated group (27/45 [60.0%] vs. 0/50 [0%], respectively) (Table 1). Representative images for the two groups are shown in Fig. 1.

**Fig.1:**
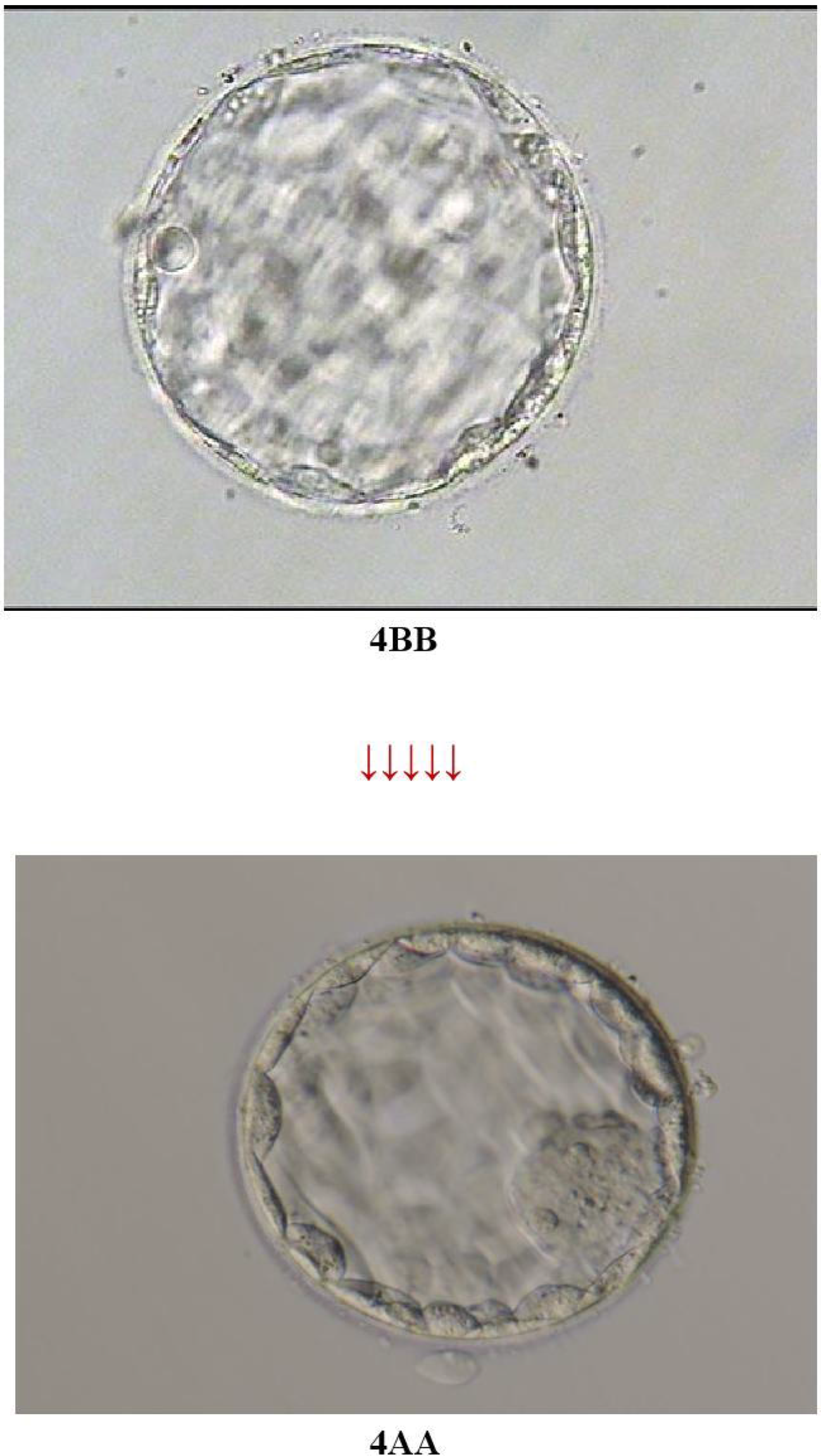
Improvements in the ICM grade and/or TE grade were observed in the ATP-exposed group (1 mM for 2 days). The representative image was shown. The grade of blastocyst before treatment was 4BB (upper image). To compare the effect of ATP treatment, we showed the representative image of same blastocyst before (upper image; 4BB) and after ATP treatment (4AA; below image).

### The mRNA expression levels of Oct3/4 and Nanog within human blastocysts

We investigated the mRNA expression levels of Oct3/4 and Nanog within human blastocysts. As a result, treatment with ATP (1 mM for 2 days) led to upregulation of Oct3/4 and Nanog mRNA expression compared with the untreated group as the control group (Oct3/4: 20.1% ± 3.3% vs. 96.3% ± 3.6% in the untreated and ATP-treated group, respectively and Nanog: 5.1% ± 3.6% vs. 35.1% ± 20.4% in the untreated and ATP-treated group, respectively). Therefore, the up-regulations of the mRNA expression levels of Oct3/4 and Nanog within human blastocysts could be observed.

## Discussion

The two manuscripts on mouse stimulus-triggered acquisition of pluripotency (STAP) cells [9, 10] were retracted in 2014. Furthermore, scientists worldwide have been unable to replicate STAP cells and/or the STAP phenomenon [11, 12].

However, investigations into STAP cells and/or the STAP phenomenon by Dr. Niwa's group at RIKEN CDB in Japan have shown that ATP can upregulate Oct3/4 and Nanog mRNA expression in mouse hepatocytes [1].

Although all human blastocysts (Day 5) that were treated with ATP (200 mM for 2 days) according to the STAP protocol [1] were bursted, the present study showed that appropriate ATP treatment (1 mM for 2 days) improved the ICM grade and TE grade in cultured human blastocysts. Although Oct3/4 is an important regulatory factor during early embryonic development [2], the ATP-induced increase in Oct3/4 expression within human blastocysts in the present study may explain the improved quality of the human blastocysts. Furthermore, Nanog is a transcription factor expressed in the ICM [3]. In the present study, Nanog expression within human blastocysts was also up-regulated by appropriate ATP treatment (1 mM for 2 days). Therefore, the ATP-induced (1 mM for 2 days) increase in Oct3/4 and Nanog mRNA expression within human blastocysts could contribute to the improved ICM grade and TE grade within the human blastocysts.

On the other hand, a study has shown that the TE grade is significantly associated with implantation and live births, whereas the ICM grade is not significantly associated with outcomes for single-blastocyst transfers [13]. Therefore, appropriate ATP treatment (1 mM for 2 days) may improve the clinical outcomes of assisted reproductive technologies practice for infertility in women. Further basic and clinical studies will be needed in the near future.

In conclusion, although ATP treatment could not replicate STAP cells and/or the STAP phenomenon by scientists worldwide [11, 12], appropriate ATP treatment (1 mM for 2 days) in cultured human blastocysts with totipotency would be helpful for infertility women.

## Acknowledgements

We are grateful to the physicians, nurses and clinical embryologists for their assistance with the design of this study and/or the experiments performed in the United States and Japan.

## Author Contributions

Concept and design: TT and MO. Performed the experiments: TT and MO. Analyzed the data: TT and MO. Writing: TT and MO.

